# Combining evolution and protein language models for an interpretable cancer driver mutation prediction with D2Deep

**DOI:** 10.1101/2023.11.17.567550

**Authors:** Konstantina Tzavella, Adrian Diaz, Catharina Olsen, Wim Vranken

**Affiliations:** Interuniversity Institute of Bioinformatics (IB2), ULB-VUB, Brussels, Belgium.; Brussels Interuniversity Genomics High Throughput Core (BRIGHTcore), Vrije Universiteit Brussel (VUB), Université Libre de Bruxelles (ULB), Laarbeeklaan 101, Brussels, 1090, Belgium.; Structural Biology Brussels, Vrije Universiteit Brussel, Brussels, Belgium.; Clinical Sciences, Research Group Genetics, Reproduction and Development (GRAD), Vrije Universiteit Brussel (VUB), Universitair Ziekenhuis Brussel (UZ Brussel), Laarbeeklaan 101, Brussels, 1090, Belgium.

**Keywords:** Protein Large Language Models, Evolutionary Information, Machine Learning, Mutation effect prediction

## Abstract

The mutations driving cancer are being increasingly exposed through tumor-specific genomic data. However, differentiating between cancer-causing driver mutations and random passenger mutations remains challenging. State-of-the-art homology-based predictors contain built-in biases and are often ill-suited to the intricacies of cancer biology. Protein Language Models have successfully addressed various biological problems but have not yet been tested on the challenging task of cancer driver mutation prediction at large scale. Additionally, they often fail to offer result interpretation, hindering their effective use in clinical settings. The AI-based D2Deep method we introduce here addresses these challenges by combining two powerful elements: i) a non-specialized protein language model that captures the makeup of all protein sequences and ii) protein-specific evolutionary information that encompasses functional requirements for a particular protein. D2Deep relies exclusively on sequence information, outperforms state-of-the-art predictors and captures intricate epistatic changes throughout the protein caused by mutations. These epistatic changes correlate with known mutations in the clinical setting and can be used for the interpretation of results. The model is trained on a balanced, somatic training set and so effectively mitigates biases related to hotspot mutations compared to state-of-the-art techniques. The versatility of D2Deep is illustrated by its performance on non-cancer mutation prediction, where most variants still lack known consequences. D2Deep predictions and confidence scores are available via https://tumorscope.be/d2deep to help with clinical interpretation and mutation prioritization.

## Introduction

The immense amount of DNA sequencing data that is now available has highlighted key similarities and differences in the biology of individuals and their predisposition for disease. Especially relevant are missense mutations in the DNA, where a codon changes to encode for a different amino acid, so resulting in random single amino acid variants (SAVs) at the protein level [1] that can be passed on by evolution [2]. The vast majority of SAVs are ‘neutral’ or ‘benign’ variants with no known detrimental effect on the function of the protein and the phenotype of an individual. Some SAVs, however, are pathogenic and cause critical protein malfunctions that have been linked to many Mendelian diseases [3] as well as predisposing to particular cancers [4],[5],[6],[7].

Our knowledge of the effect of SAVs is still limited, with only ∼2% of the more than 4 million observed missense variants clinically categorized as either pathogenic or benign [8]. For complex diseases such as cancer, the interpretation of the impact of SAVs is even more challenging. The behavior of cancer cells cannot be attributed to a single cellular process, but is rather the result of stochastic, somatic mutations that accrue over time, with additional contributions from genetic (germline) and epigenetic variation [9]. Among these SAVs, a small subset alters the cellular phenotype, and an even smaller subset confers a selective fitness advantage to clones, enabling them to evade the stringent cell control mechanisms that would normally result in cell death via apoptosis. These mutations, known as driver mutations, play a critical role in cancer development and progression and their identification is essential for both understanding the molecular mechanisms of cancer and for developing targeted therapies [10]. Pan-cancer genome analyses have revealed that, on average, there are 4 to 5 driver mutations per cancer sample in both coding and non-coding genomic regions. However, from the sequenced samples collected from individuals with cancer, 5% of cases do not contain any of the known driver mutations [11]. In the many samples with known driver mutations, on the other hand, the possibility remains that other driver mutations were overlooked.

Given the sheer number of SAVs and the cost of experimental testing, Artificial intelligence (AI) approaches have rapidly become key tools in their putative classification. Commonly used AI approaches to identify the phenotypic effect of single (or multiple) SAVs such as SIFT [12], PolyPhen2 [13], CADD [14] and FATHMM [15] typically consider the conservation of amino acid in single sequence positions to predict the effects of mutations. However, models that capture specific pairwise epistatic interactions between amino acids, where one amino acid change requires compensatory adjustments in other parts of the sequence to maintain protein function, have been shown to enhance mutation effect prediction [16],[17],[18],[19]. Addressing higher-order epistasis, involving more than two amino acids in different positions in the sequence, remains a challenge despite numerous converging evidence on their crucial role in protein evolution and fitness [20],[21],[22]. To our knowledge, only EVE [23] and AlphaMissense [8] are able to incorporate such higher-order constraints. EVE relies on evolutionary information learned by amino acid sequences for each protein across different species using Multiple Sequence Alignments (MSAs) and clusters the learnt representations into benign or pathogenic groups based on their likelihood to occur. Because alternative isoforms of the same gene have identical homologs, it remains uncertain whether this approach can differentiate the effect of mutations on different isoforms [24]. Also, as an unsupervised model, it encounters challenges to discern protein-specific structural or functional alterations. To capture protein-specific nuances, the integration of direct functional and contextual data for a particular protein has been shown to lead to substantial improvements in prediction [25],[26]. AlphaMissense incorporates such data in a supervised setting, but considers frequently observed variants in human and primate populations as benign, whereas absent variants are labelled as pathogenic, resulting in an inherently noisy and biased training set, as rare variants can also be benign [27]. None of these predictors provide an interpretation of results that could support possible clinical decisions based on them. Another deep-learning method for variant effect prediction utilizes protein language models, a technique originating from natural language processing. Protein language models (pLMs) do not rely on explicit homology and can estimate the likelihood of any possible amino acid sequence, learning from common patterns across protein families, allowing information to be generalized [28], [29], [30]. They have demonstrated the ability to implicitly learn how protein sequences influence various aspects of protein structure and function, including secondary structure and subcellular location [31], [32]. However, they have not yet been tested on the challenging task of cancer driver mutation prediction at a large scale.

We present D2Deep, a protein sequence-only prediction method to distinguish driver from passenger mutations in cancer that is based on an original combination of protein-specific evolutionary information (EI) from MSAs with pLMs. PLMs inherently represent the probability of amino acids occurring in various contexts, in other words the grammar (or syntax) of natural protein sequences, while MSAs represent protein-specific evolutionary information that incorporates the effect of each mutation on protein function (semantic correctness). By employing a Gaussian Mixture Model (GMM) on the pLM features across evolutionary related sequences, we can therefore capture the site-specific evolutionary variation of proteins and so detect when sequences are semantically incorrect. The features extracted from this approach, which now combine general protein sequence principles with specific protein information, are subsequently used in a downstream supervised task for distinguishing between driver and passenger mutations in cancer. They capture the effect of the mutation throughout the protein and can be used for the interpretation of results in the clinical setting. To minimize bias, we meticulously curated a per-gene balanced training set of somatic mutations, and trained a generic multilayer perceptron (MLP) predictor that can score pathogenicity for any gene variant (Fig.1). Importantly, the statistical nature of the model allows for the confidence calculation score, relying on its fit on the evolutionary data. We validate this model on an independent test set, built to eliminate bias of cancer related genes, and performs to state-of-the-art standards whilst only using generic protein information. Its general applicability is further demonstrated in non-cancer mutation prediction, where still over 98% of variants lack known consequences.

**Fig. 1.**
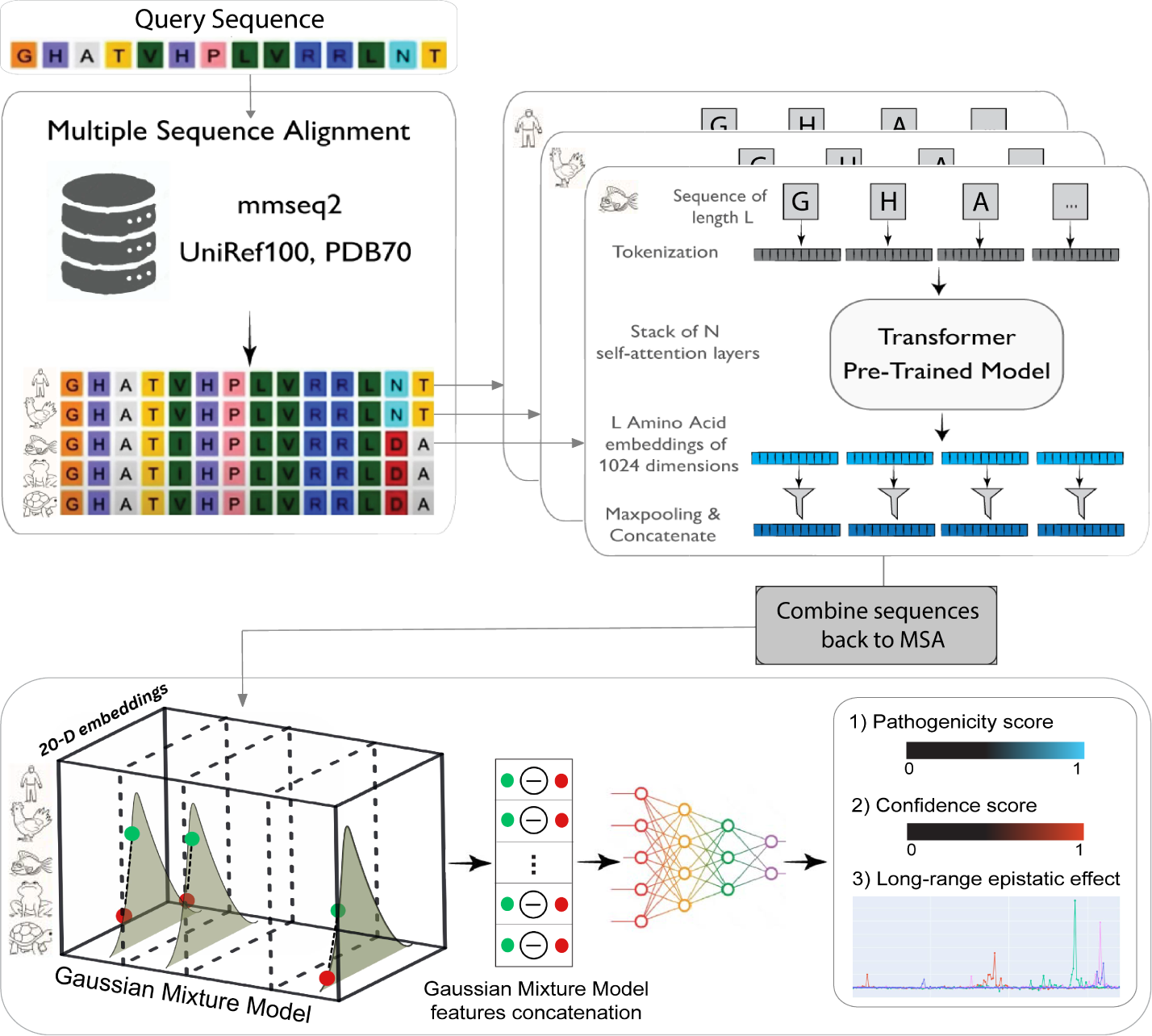
Overview of D2Deep’s pipeline. For the mutated protein sequence, a genetic database search is performed for the production of Multiple Sequence Alignment from various species. Each Amino Acid of the MSA is mapped to the 20-D pLM representations vector. The pLM vectors are combined back to the MSA and a Gaussian Mixture Model is fitted to the representations of all species on each MSA position. The GMM log-probability of the Mutant’s representation is then subtracted from the GMM log-probability of the Wild-Type’s representation. The differences are subsequently concatenated to capture the effect of the mutation over all positions of the sequence and used as features for the Fully-Connected Neural Network. For simplicity, a 1-dimension GMM is shown on the Figure.

## Results

### D2Deep performance compared to state-of-the-art

The training data for cancer driver genes is significantly biased, with very frequently studied cancer genes much more prevalent in these datasets, resulting in predictors often not interpreting the mutation itself, but rather deciding on whether the mutation occurs in a well-known cancer related gene [2]. We thus put strong emphasis on training D2Deep on cancer balanced data. The dataset consists only of mutations in Cancer Gene Census oncogenes or Tumor Suppressor Genes, so avoiding reliance on prior knowledge about the gene being a known driver gene or not (see Methods). In the D2Deep approach, the mutated sequence’s MSA is processed by mapping each amino acid to a 20-dimensional pLM representation vector. Subsequently, a GMM is employed for each position, calculating the differences in log-probabilities between the Wild-Type and Mutant, thereby capturing the mutation’s impact across the entire sequence (Fig.1). To evaluate D2Deep’s performance in classifying passenger from driver mutations, we calculated the predictions on the cancer specific, unbiased test set DRGN and compared its performance to 6 state-of-the-art predictors SIFT [12], DEOGEN2 [33], EVE [23], FATHMM-cancer [15], CADD [14], PolyPhen2 [34] and AlphaMissense [8]. Our method outperforms all but the FATHMM-cancer predictor as shown on Fig.2a-d. FATHMM-cancer’s efficacy stems from its specialized training on the cancer mutation frequencies within the conserved domains of each protein. To investigate if FATHMM contains an inherent bias due to its training set, we segregated the mutation into the known hotspot cancer mutations and non-hotspot ones. Indeed, as shown in Fig.3a, the performance of FATHMM drops significantly between hotspot and non-hotspot mutations. AlphaMissense, with the third highest performance, has the worst performance for non-hotspot mutations while D2Deep consistently performs well on the hotspot and non-hotspot mutations. Because reliable annotations for cancer related mutations are scarce, a part of the DRGN set is also included in our Training set (Supplementary Fig.7). To study how the inclusion of test mutations in the training set affects the performance of the algorithm we used cross-validation (5-folds) where the test set of each fold was excluded from the training set. Again D2Deep has comparable performance to the state-of-the-art (Fig.2b). Here we have to note that the rest of the predictors were not re-trained to exclude the test mutations and thus D2Deep is more strictly tested than the others. As the predictors do not contain all protein isoforms of the test set, we show both the overlapping mutations present in all predictors (1868 mutations) and when the predictors are penalized for missing values (3603 mutations).

**Fig. 2.**
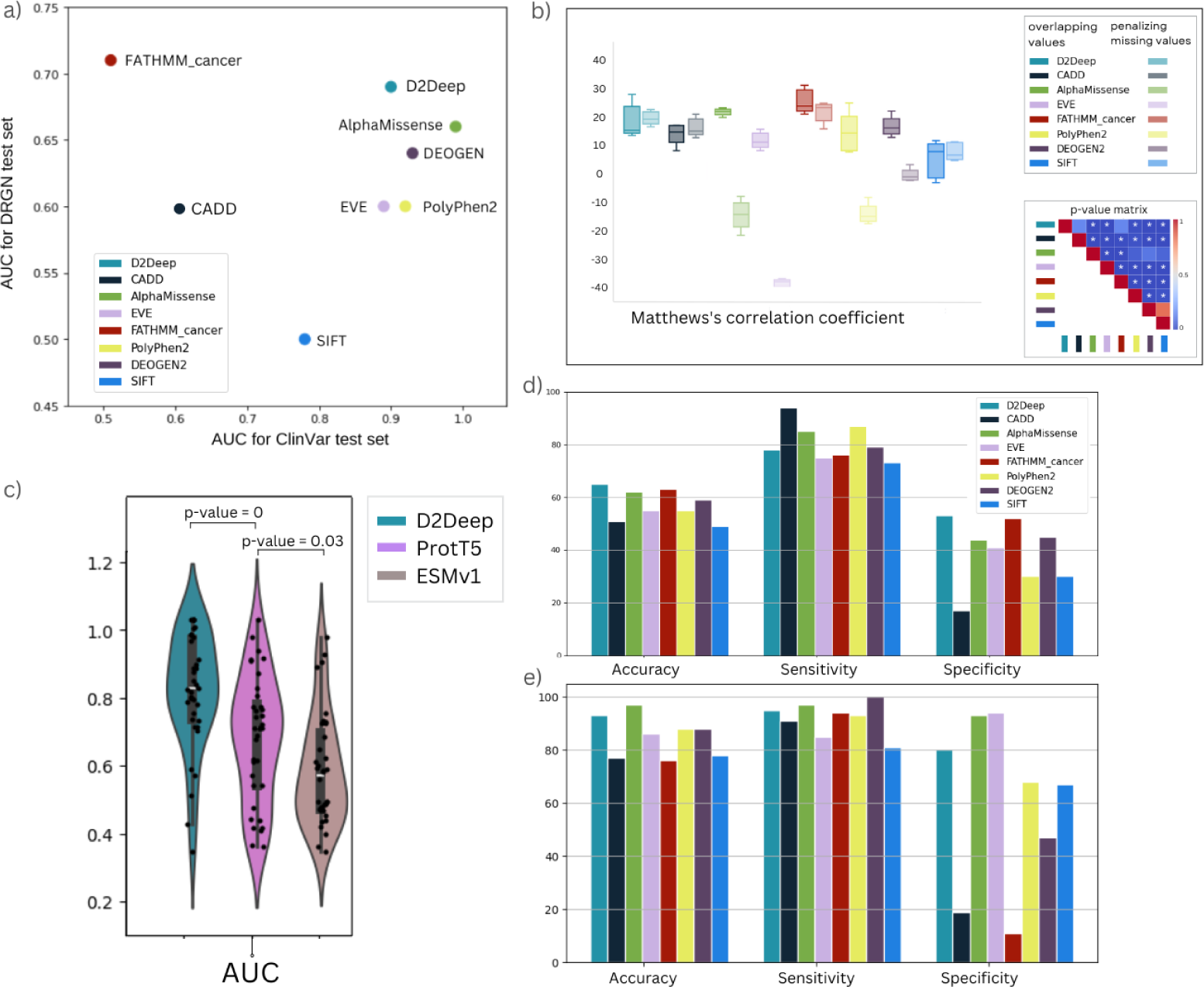
D2Deep compared to state-of-the-art. a) Comparison of all six predictors for both ClinVar and DRGN test sets based on AUC metric -b) 5-fold validation Matthew’s correlation coefficient of predictors on DRGN variants for overlapping values (intense colors) and when missing values are penalized (soft colors), significant p-values (*<* 0.05) are marked with asterisks in p-value matrix - c) Leave one out cross validation using D2Deep, ProtT5 and ESM-1v features and subsequently inputted into a supervised model - d) D2Deep results for DRGN test set compared to existing predictors - e) D2Deep results for ClinVar annotations on cancer driver genes compared to existing predictors.

**Fig. 3.**
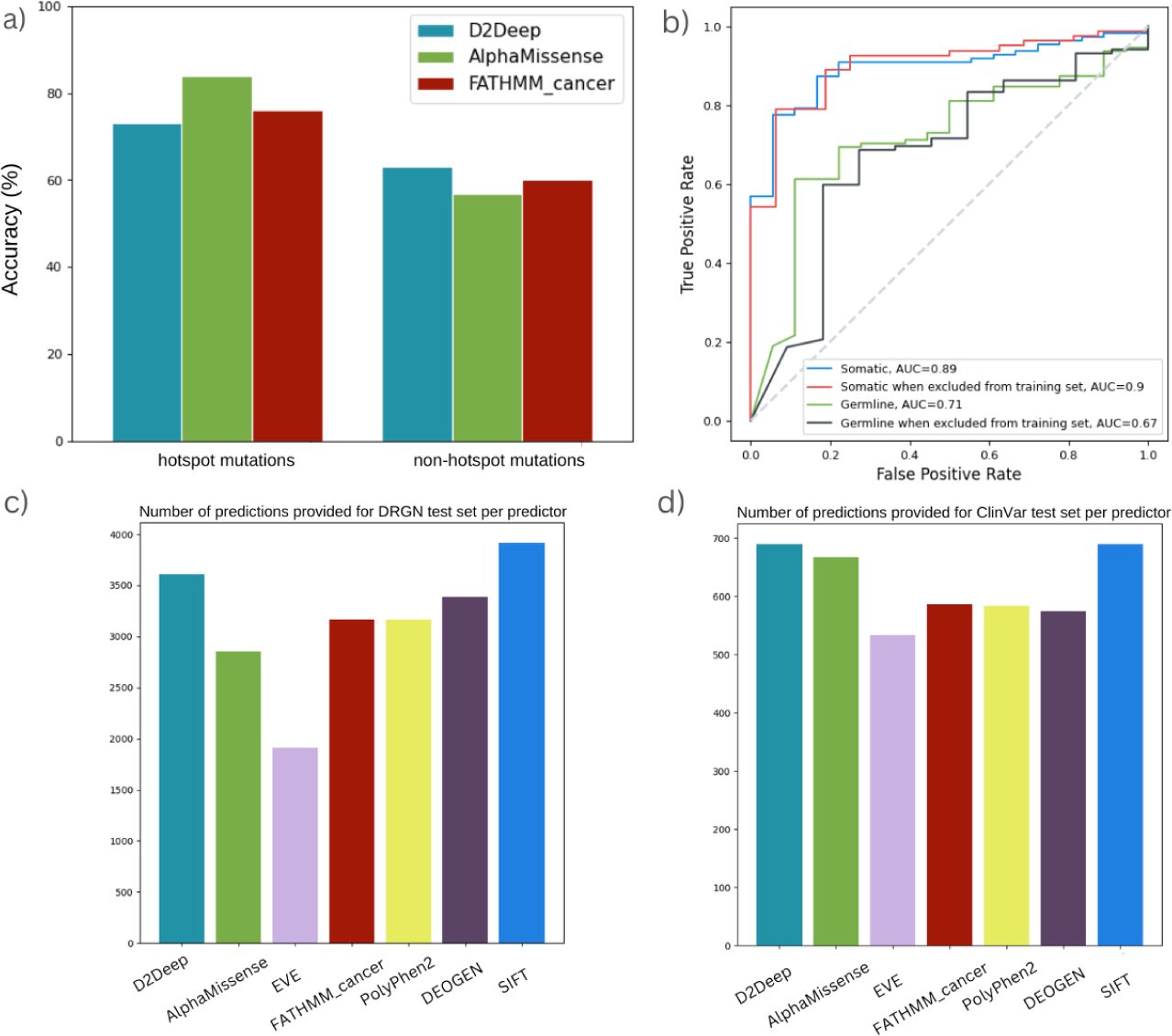
a) Performance of AlphaMissense and D2Deep on hotspot cancer mutations compared to mutations not on hotspot regions - b) Comparison of D2Deep’s performance on somatic versus germline ClinVar mutations. Somatic have higher performance that the germline ones and without drop in performance when excluded from the training set - c-d) Number of valid predictions for DRGN and ClinVar test sets respectively for all predictors.

We further tested D2Deep’s ability to predict fitness across cancer gene variants. To this end, we benchmarked D2Deep on 690 confident labels from ClinVar, for which there are no conflicting annotations, for five cancer driver genes used in clinical setting. Here again D2Deep is comparable to state-of-the-art predictors while not trained on ClinVar data, as shown on Fig.2e. Overall, D2Deep consistently excels in both test sets (Fig.2a), proving its effectiveness and ability to generalize using only sequence information. In contrast, the highest-scoring predictors employ a range of contextual annotations. AlphaMissense, for example, relies on Structural context by using an AF-derived system. Polyphen-2 is trained on binding and functional site annotations as well as the output of other predictors. DEOGEN2, on the other hand, uses a variety of contextual information such as gene and pathway-oriented features.

Finally, D2Deep was tested on a dataset that includes only somatic and germline ClinVar mutations without conflicted annotations. The same amount of pathogenic and benign mutations were retained per protein for the germline and the somatic group, leading to a test set of 129 mutations for each. The somatic performance was higher than for the germline mutations (Fig.3b) as expected due to the somatic-specific training set used. Accordingly, excluding the germline mutations from the training set (resulting in 113 test mutations) led to a slight decrease in performance, not observed for the somatic ones.

Most human genes undergo alternative splicing, which means that a single genetic variant can have different effects on various protein isoforms. D2Deep’s only requirement in sequence provides an advantage for prediction of different isoforms of the same protein and Variants of Unknown Significance compared to other predictors, as shown in the number of predicted mutations in DRGN and ClinVar tests sets respectively (Fig.3c-d).

We provide the pre-calculated predictions for all possible mutations for the Compermed cancer gene panel (see Methods) on our webserver https://tumorscope.be/d2deep (Supplementary Fig. 8)

### Evolutionary Information aids Large Language models in capturing the specific effect of mutations

Language models for proteins are trained on extensive sequence databases such as those found in UniProt [35] and Pfam [36]. As a result, the representations they acquire primarily capture general contextual information applicable to a wide range of proteins, rather than being specifically tailored to the protein under study. To probe how much the protein-specific evolutionary information from the MSAs used in D2Deep adds to the features directly captured by the pLM model, we tested three sets of features as inputs to the same predictive model (see Methods). We compared the D2Deep features to the pLM features extracted from ProtT5 [31] and from ESM-1v, a pLM specifically trained on mutations [37]. We first tested the three approaches in a leave one out cross-validation setting on the balanced, somatic dataset. The beneficial effect on model performance obtained by i) introducing information from the MSA and ii) extracting features through our GMM-based approach is then evident from Fig.2c, We also tested D2Deep features using only the pLM-based features in the unbiased, independent DRGN test set (gray points, Supplementary Fig.9) and compared it with the model that integrates both pLM-based and EI features on a 5-fold cross-validation setting. Again the inclusion of EI increases the performance as shown in the coloured points in Supplementary Fig.9. Plotting the features of DRGN mutations on four random cancer genes in a 2-D space demonstrates that the pLM features fail to capture mutation-induced changes but effectively represent protein-family information, as evident in the protein-specific clustering (Supplementary Fig.10). Therefore, incorporating EI from the MSA provides additional mutation-specific semantics, beyond the general protein syntax captured by the pLM.

### Certainty and Confidence

In a clinical context such as cancer it is crucial to be able to assess the reliability of a prediction. We therefore explored whether we could use the log-likelihood measure inherent to GMMs, which quantitatively assesses the goodness of fit of a data point, as a proxy to reliability. A higher log-likelihood indicates a better fit, suggesting that the GMM effectively captures the underlying structure of the data. We calculated the confidence score equation using grid search as described in Methods. The weighted accuracy on the DRGN test set was increased from 64.2% to 65.3% and the high confidence mutations for genes TP53, AR, PTEN, CHEK2 and BRAF have increased performance as shown in Table 1. In Fig.4(column b) and Supplementary Fig.11 we show the qualitative performance when only the most confident predictions (with a threshold of 65%) are kept.

**Fig. 4.**
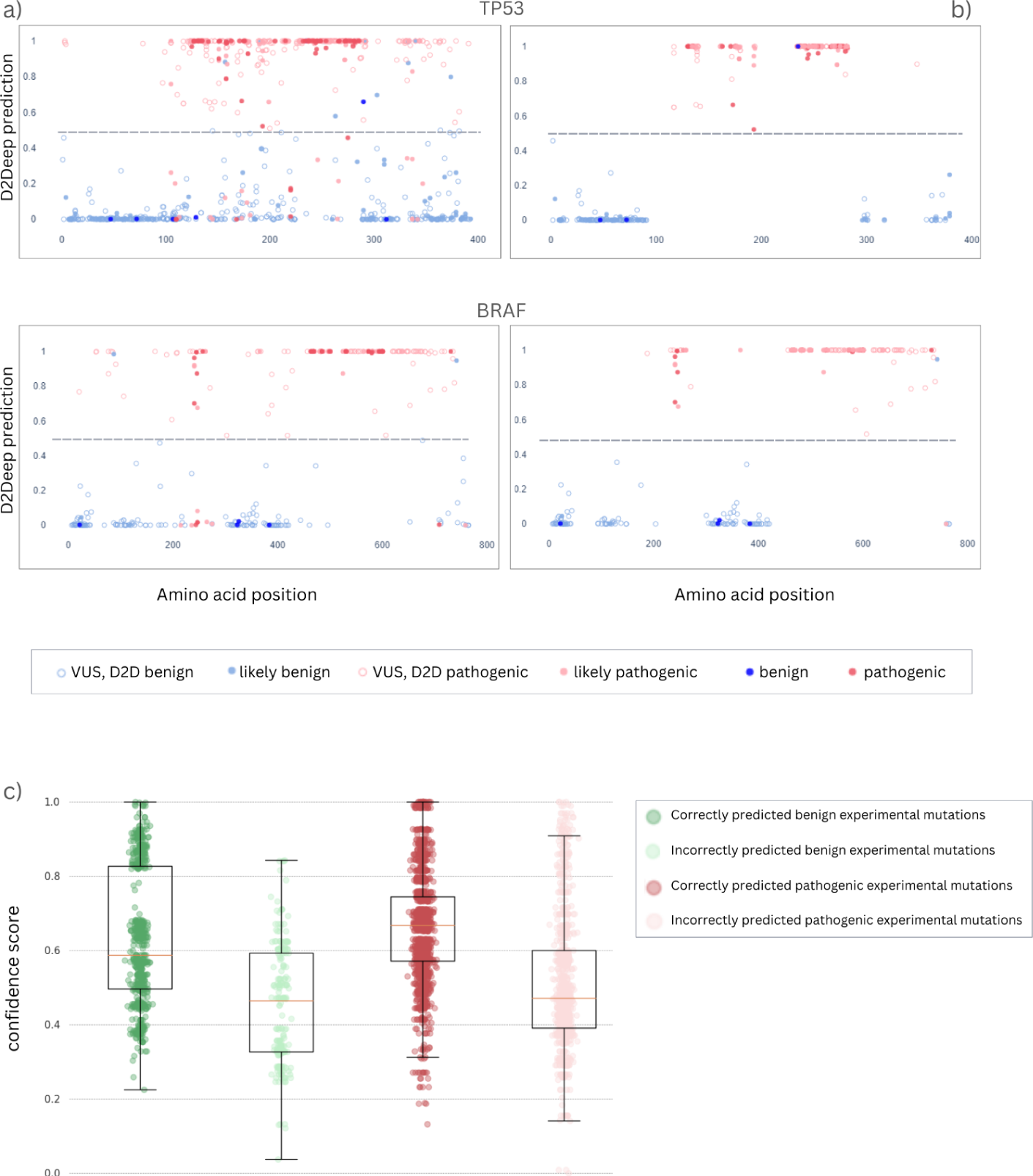
D2Deep confidence scores. Predictions for ClinVar annotations on 2 cancer driver genes (TP53, BRAF) when a) all D2Deeps’ predictions are shown - b) high confidence predictions for Clin- Var annotations are shown - c) D2Deep confidence scores of correctly (intense colors) and incorrectly (soft colors) classified mutations for TP53 gene. We chose to demonstrate the predictions of genes with a relatively large number of high-quality labels in ClinVar. The rest of available predictions can be found on our webserver.

**Table 1.** D2Deep performance on ClinVar annotations.

| Gene | ClinVar annotations | Accuracy (%) | Weighted Accuracy (%) | VUS annotations |
| --- | --- | --- | --- | --- |
| TP53 | 280 | 85 | 91.5 | 627 |
| BRAF | 109 | 89 | 96.9 | 243 |
| AR | 116 | 86.2 | 93.5 | 67 |
| CHEK2 | 10 | 90 | 94.2 | 1498 |
| PTEN | 175 | 81.7 | 96.1 | 586 |

Increase of D2Deep’s performance for known cancer driver genes TP53, BRAF, AR, CHEK2 and PTEN when confidence score is used.

We tested D2Deep on independent Deep Mutational Scanning data [38], where all mutations in all protein positions are tested in relation to an experimental fitness measure such as cell function. Advantageous mutations have positive Enrichment Ratio score (ER) while deleterious for the function of the cell have negative ER. The study also includes synonymous (neutral) mutations with scores near zero, which were excluded from the analysis to avoid potential ambiguity. Accordingly, an exclusion range was defined around zero (-0.2 ≤ ER ≤0.2). Mutations with scores exceeding 0.2 were classified as benign, while those with scores less than -0.2 were classified as pathogenic. By design, D2Deep predictions have a lower value (≤ 0.5) for benign and a higher (*>* 0.5) value for pathogenic mutations. As a consequence, in the case of TP53 gene, a negative correlation with the ER scores is observed with a Pearson correlation of -0.46. Furthermore, we selected the confidently benign experimental mutations (ER score ≥ 0.2) and grouped them based on D2Deep predictions as benign or pathogenic. We did the same for the confidently pathogenic experimental mutations (ER score ≤ -0.2) that were correctly and incorrectly classified by D2Deep. The correctly predicted mutations have significantly higher prediction confidence scores than those incorrectly predicted in both groups, with Wilcoxon rank sum p-value equal to 1.52e-11. (Fig.4c).

### Interpretation of epistatic features of known and novel mutations

There are few predictors that capture specific epistatic interactions between amino acids. D2Deep does so by capturing changes inflicted throughout the sequence when a mutation is introduced. To illustrate this behaviour, we first compared the D2Deep features with long-range effects proposed by MAVISp [39] for the KRAS gene (Fig. 5). MAVISp uses the protein’s structure to calculate the free energy changes caused by the mutations on the Switch I region, suggesting that some of these changes might contribute to protein activation through distal effects. In accordance with MAVISp findings, the D2Deep features have higher signal at positions corresponding to peaks in allosteric free energy changes, relying solely on sequence information.

**Fig. 5.**
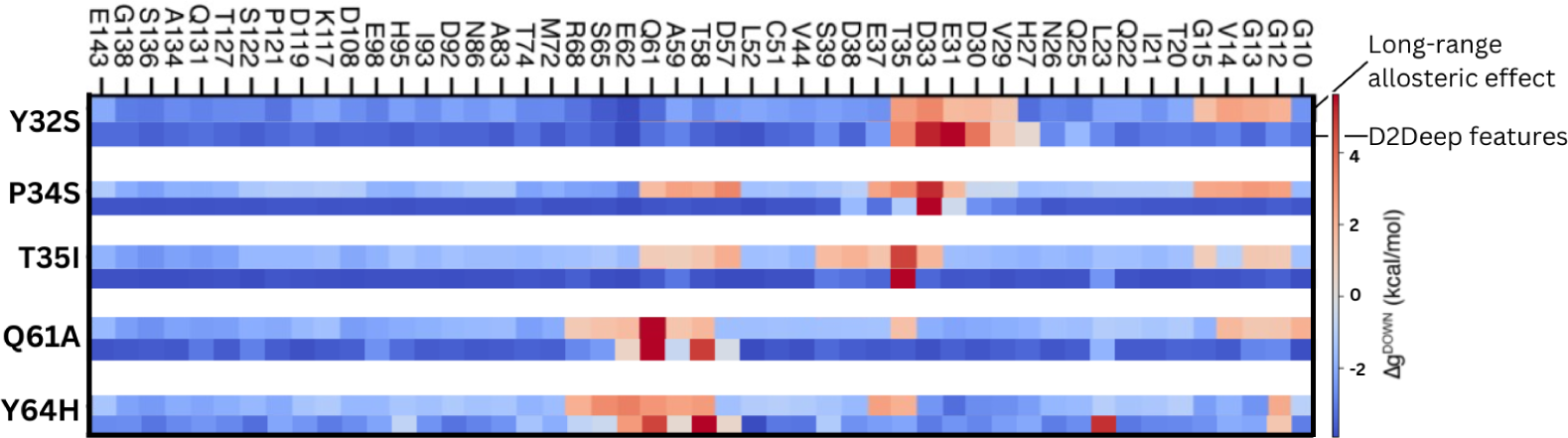
Long-range effect of KRAS mutations. Variations in free energy calculated by MAVISp caused by KRAS mutations on y-axis (’Long-range allosteric effect’). On the heatmap blue denotes stabilization through reduced dynamics, while red signifies increased motion, indicating a destabilizing effect. The mutations potentially influence the distant sites (at least 8 Å from the original mutation), displayed on the x-axis. On the second rows of each mutation, we show the D2Deep features for the same distal sites.

Additionally, we collected ClinVar benign and pathogenic mutations on the TP53 gene and analyzed each mutation’s impact on distant amino acids. Fig.6a shows the features of two pathogenic mutations, E258K and C238R. Their features have the highest values, and thus influence, distant in sequence amino acids. Nevertheless, when depicting the AlphaFold2 predicted structure (Fig.6b), it becomes apparent that the peaks of these features are, in reality, indicative of nearby amino acids in three-dimensional space. This suggests that D2Deep features can capture long-range interactions between amino acids.

**Fig. 6.**
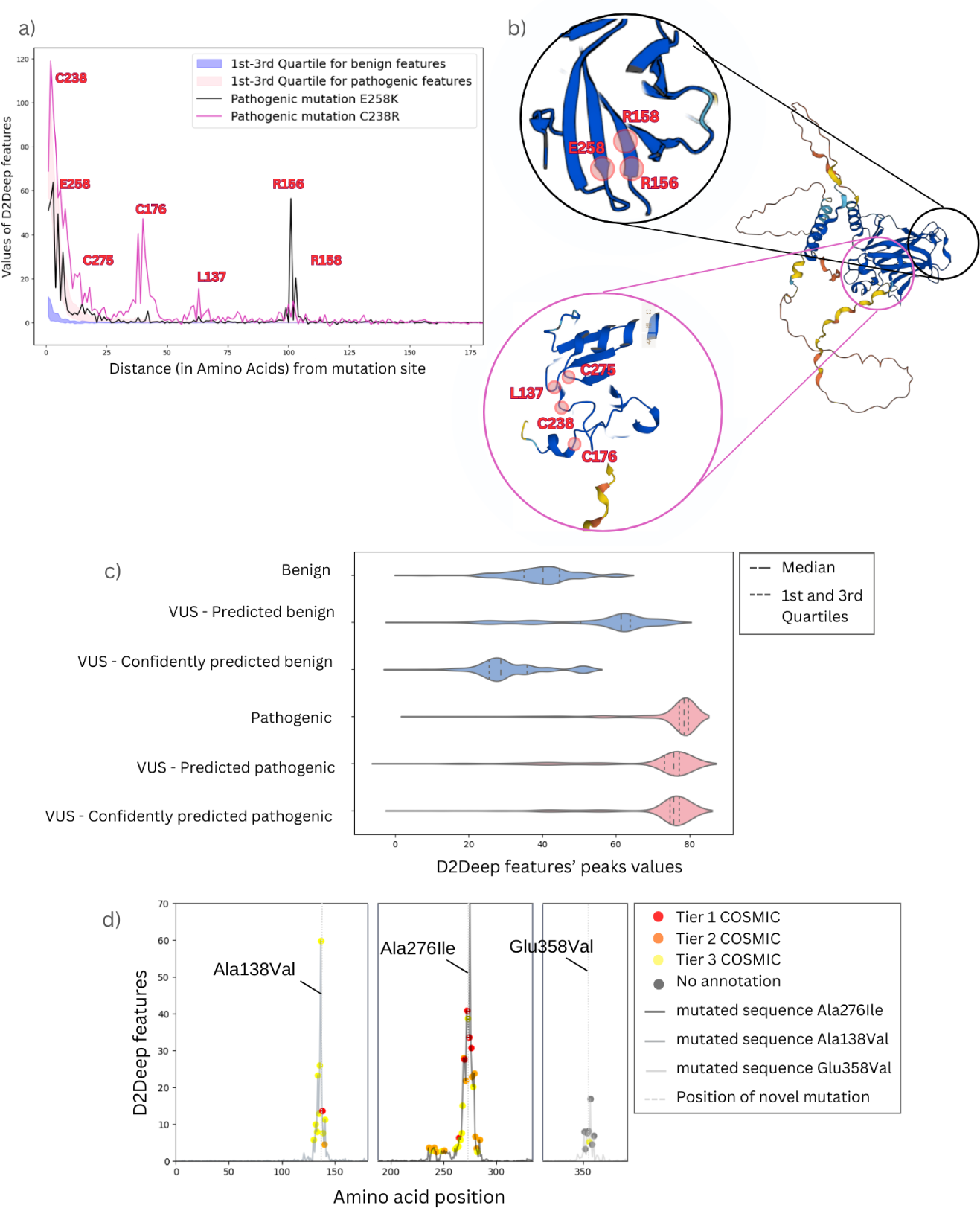
Long-range effect of TP53 mutations. a) D2Deep features used in the downstream prediction of 149 benign and pathogenic ClinVar TP53 mutations. Two pathogenic mutations E258K and C238R were considered to study the influence of features in distant amino acid and capture potential co-evolution forces - b) Predicted AlphaFold2 TP53 structure structure with focus on the surrounding regions of the mutations E258K and C238R. The amino acids that correspond to the highest D2Deep features are highlighted - c) D2Deep features for benign (blue) and pathogenic (red) mutations of TP53. ClinVar benign mutations (352 mutations) have low feature signal while ClinVar pathogenic mutations (176 mutations) have high D2Deep signal. The VUS features follow the ground truths suggesting a correlation between known and unknown mutations. We consider the top ten highest values as peaks - d) D2Deep features of novel mutations Ala138Val, Ala276Ile and Glu358Val. Only the sequence region where the D2Deep features are most affected by the mutation are shown, with the most influenced amino acids enriched in COSMIC mutations.

To further investigate this, we tested the features of novel somatic driver mutations on the TP53 driver gene from three recent publications [40], [41], [42]. These mutations have been experimentally validated. Two of them (Ala138Val and Ala276Ile) are predicted by D2Deep as pathogenic, with confidence 69% and 90% respectively, while Glu358Val is predicted as benign, with 50% confidence. These results were consistent with those obtained from state-of-the-art predictors. We again depicted the features of mutated sequences Ala138Val, Ala276Ile and Glu358Val. The features exhibit elevated values in positions coinciding with established driver mutations sourced from the COSMIC database (Fig.6d). This suggests that D2Deep features can associate novel mutations with confirmed ones.

When only considering the magnitude of D2Deep features, TP53 mutations labeled VUS in ClinVar that are predicted as benign by D2Deep have a lower feature signal, in accordance with the feature range of confirmed benign labels taken from ClinVar Fig.6c. The same is also true for the confirmed ClinVar pathogenic mutations, with the tendency reinforced when we consider confident (score ≥ 50%) predictions. Correlating ground truth signals with predictions in this way should help in the interpretation of VUS in the clinical setting, where mutants that connect within the feature space to known driver mutations could be prioritised. Furthermore, the effect of a novel mutation is more confidently recognised by the Neural Network when known pathogenic mutations are present, which likely give similar feature signals to the novel mutation (Supplementary Fig.12).

## Discussion

Distinguishing between passenger and driver mutations in cancer is a complex problem that can only be tackled by judicious use and interpretation of data. In D2Deep, we achieve this by combining general protein sequence semantics from language models with protein-specific evolutionary information.

D2Deep models the impact of a mutation throughout the sequence, so better capturing the intricate mechanisms of epistasis whilst only using protein sequence based information. Its power is illustrated on cancer-specific and non-cancer specific test sets, demonstrating that a predictor that utilises only sequence information can in this way consistently outperform state-of-the-art predictors trained with gene, pathway and domain-specific information. Consequently, it can provide valid predictions for sequences lacking fixed structure or domain annotations. Most notably, our method offers a quantifiable measure of the mutation’s impact on the protein, allowing for meaningful comparisons between known and unknown mutations. Variants of Uncertain Significance, which continue to pose challenges for scientists and clinicians, can subsequently be prioritized according to the predictions in combination with experimental identification of novel driver mutations. An added advantage of D2Deep is its statistical nature, which makes it possible to calculate a confidence score for each prediction, enabling better prioritization of novel mutations and setting it apart from other pathogenicity predictors. All predictions and data are available at https://tumorscope.be/d2deep, which will be regularly updated with new genes.

A shortcoming of this method is its reliance on ProtT5, limiting input sequences to fewer than 2200 amino acids. We hope future improvements will address this to cover all cancer-related genes. Other pertinent challenges not addressed by this work are, first and foremost, the differentiation of the mutation’s impact depending on the phenotype. It is widely recognized that mutations within the same gene, as well as identical variants, can result in varying degrees of disease, and even give rise to distinct disorders. The inclusion of cancer types or other histopathological information will therefore become necessary for even more precise predictions. In addition, we hope to extend the D2Deep capabilities to also suggest which molecular phenotype changes, such as increased activity or binding, are responsible for the driver categorisation of a mutant. Finally, driver mutations can remain dormant for extended periods and may initiate oncogenesis only combined with other mutations. Multiple mutations on the same gene have not been considered for this work, but the method is inherently capable of capturing these. In the future, our approach can also be synergistically integrated with supplementary contextual information, such as cancer type, and harness the power of sequence-only features to address combinations of variants, leading to enhanced predictions that will aid clinicians in identifying various cancer subtypes and ultimately facilitate more effective treatments, even when considering diseases beyond cancer.

## Methods and Materials

### Training set

We curated 3,093,303 SAVs from the Catalogue of Somatic Mutations in Cancer (COSMIC - Cancer Mutation Census releasev92) [43]. We excluded pseudogenes and RNA genes and kept 3825 mutations that were annotated as Pathogenic or Likely Pathogenic in cancer-related diseases in ClinVar and are recurrent in a Cancer Gene Census oncogene or in a Tumor Suppressor Gene (namely Tiers 1, 2 and 3 in COSMIC as described in Supplementary Fig.13). Their distribution in cancer-related diseases in ClinVar can be seen in Supplementary Fig.14. We additionally collected 2,657 somatic variants with known molecular consequences from Cancer Genome Interpreter (CGI). To balance the (likely) pathogenic SAVs of each gene with a balanced benign set, we curated the UniProtKB/Swiss-Prot humsavar dataset (release 21st December 2021) keeping the 39,325 Benign/Likely Benign SAVs. To this we added the ClinVar dataset with benign annotations that resulted in 43,030 benign SAVs. Additionally we included common variants, SAVs frequently observed in the general population, from the gnomAD database, keeping mutations with Allele Frequencies (AF) higher than 0.1%. Lastly, we mined single nucleotide polymorphisms (SNPs) from the Single Nucleotide Polymorphism database (dbSNP) excluding mutations with conflicted or uncertain interpretation, pathogenic or risk associations. The mapping of the chromosomal position provided by dbSNP to protein coding positions was performed with the use of TransVar software [44]. After removing dataset overlap, 178,979 benign SAVs were retained. To establish a well-balanced training set at the gene level, we gathered both pathogenic and benign mutations for each gene while maintaining a ratio of not exceeding the 40-60% class balance. To ensure fairness, for genes with a limited number of available mutations, we enforced a maximum difference of two mutations between the pathogenic and benign classes. After the filtering out the sequences with more than 2200 amino acids, the final training set contained 6,608 mutations, 2,956 deleterious and 3,652 benign, internally balanced within 1,012 genes. The script and data for the above workflow are publicly available.

### Test sets

**DRGN**: To assess the construction bias affecting the in-silico predictors, researchers introduced the DRGN set [2], comprising a total of 4,093 variants. Among these variants, 1,809 were identified as deleterious, while 2,284 were categorized as passenger mutations. These variants were mapped to 153 driver genes. The deleterious variants were specifically selected from CGI, based on their experimental validation as cancer driver variants [45]. As for the passenger mutations, they encompassed 63,525 germline variants unrelated to cancer, sourced from Humsavar [35]. To ensure a comprehensive evaluation, the final test set included genes that had at least one positive (deleterious) and one negative (passenger) sample, comprising 3608 mutations.

**Consensus Pathogenic Variants**: We calculated the output of the method on 269 cancer driver genes included in Next Generation Sequencing (NGS) panel of biopsies of haematological and solid tumours from Compermed Guidelines (https://www.compermed.be/docs/Guidelines jan2020 2.pdf). For the demonstration of predictions, we chose genes with a relatively large number of high-quality labels in ClinVar, three Tumor Suppressor Genes (TP53, PTEN, CHEK2) and two oncogenes (BRAF, AR) with Review status: Practice guideline, Expert panel, Multiple submitters, Single submitter (downloaded in March 2023).

### Transfer learning with Pre-Trained model ProtT5-XL

Language models (LMs) are typically based on the Transformer architecture, which uses a mechanism called ‘self-attention’ to weigh the importance of different words in a sentence when making predictions. This allows them to capture long-range dependencies in text. Protein LMs (pLMs) leverage vast datasets of protein sequences, similar to the training of language models. By training on billions of amino acids, these models learn to model statistical dependencies among amino acids based on their cooccurrence patterns across sequences. The learnt representations have been shown to retain critical biophysical properties, making them valuable inputs for downstream tasks [31]. The best-performing pLM for our task was ProtT5-XL [31], trained on the BFD100 [46] and Uniref50 [47] datasets, from which we extracted 1024-dimensional (1024-D) amino acid representations.

### Gaussian Mixture Model

The Multiple Sequence Alignments (MSAs) for both the training and test sets were generated using the mmseq2 algorithm with the Uniref100 and PDB70 databases [47]. The resulting alignments were filtered according to the protocol proposed by Hopf et al. [15]. In line with this protocol, we retained sequences that aligned to at least 50% of the target sequence and had at least 70% column occupancy. For five protein isoforms in our training set (out of a total of 1,132), the filtering process resulted in fewer than two aligned sequences in the MSA. To address this, we applied a more lenient filtering cutoff, retaining sequences with at least 20% identity to the original protein.

Each amino acid in the MSA was then mapped to the 1024-dimensional (1024-D) embedding learnt during the pre-training phase. We applied a Gaussian Mixture Model (GMM) from the sklearn library to each MSA column to measure how mutations deviate from the underlying distribution. Since the feature space (1024-D) exceeded the number of sequences in some MSAs, which could lead to overfitting, we performed dimensionality reduction. We reduced the feature dimensions using max-pooling with a kernel size of 50 and a stride of 0, resulting in 20-dimensional vectors (20-D) for each amino acid. The selection of 20-D was based on an evaluation of different dimensions to determine which provided the highest probability for the GMM and thus the best fit.

To prevent bias from the GMM’s structure in each dimension, we employed a threshold-based approach. We fitted a GMM to the 20-D samples of each MSA column, computed the log-likelihood of the WT features to be in the model and the log-likelihood threshold, below which 1% of the samples lie. Then, we computed the distance between the log-likelihood and this threshold (WT-threshold). When a mutation is introduced, the pre-trained embeddings of the entire sequence are affected. To capture its effect, we calculated the distance between the mutation’s log-likelihood and above 1% threshold (MUT-threshold). This allowed us to quantify the mutation’s impact, even when the WT and mutation features were similar in magnitude but opposite in sign.

The final feature was obtained by computing the difference between WT-threshold and MUT-threshold at each sequence position, yielding one value per position. These differences were concatenated across the sequence to form the feature set used in the downstream supervised learning task.

### Model Architecture and Training

We built a deep learning model for the mutation classification. The model receives the features produced by the GMM part of the algorithm and predicts the mutation’s pathogenicity as a probability value from 0 to 1. The classifier is composed of two Fully Connected (FC) layers followed by a batch normalisation for faster and more stable training. Because the FC layers must have a defined input length, we chose as maximum length the 2200 amino acids, padding shorter sequences with zero, which did not affect 90% of the cancer gene panel (Supplementary Fig.15). To select the hyperparameters of D2Deep, we performed a grid search using 90% of the training data to train the model and the remaining 10% as the validation set to select the hyperparameters. The test sets were not used for hyperparameter selection. The model was trained for 200 epochs. The number of epochs to train were selected based on the early stopping technique choosing the epoch on each the validation error starting increasing while the training error continue decreasing. Dropout layers of 0.3 are applied during training to all layers to avoid overfitting, followed by a Rectified Linear Unit (ReLU) activation function. The final FC layer is used to decide pathogenicity with the use of a sigmoid activation function incorporated in the BCEWithLogitsLoss loss function. The use of BCEWithLogitsLoss is recommended in the PyTorch documentation (https://pytorch.org/docs/stable/generated/torch.nn.BCEWithLogitsLoss.html), over a plain Sigmoid followed by a BCELoss, as numerically more stable). The weights of the classifier were initialised using Xavier normal initialization and the batch size was set to 64. Instead of depending only on the current gradient to update the weights in every step, a gradient descent with momentum of 0.1 aggregating current and past gradients. The optimizer used was AdamW with a learning rate of 0.00003 and a scheduler with a warmup period of 1000 steps. The model was trained in the PyTorch framework.

### pLM comparison

For the comparison between pLMs and pLMs and EI, we follow the workflow proposed by ESM-1v [37]. We first calculated the pLM embeddings for the training dataset using the EMS-1v (esm1v t33 650M UR90S 1.ptk) pLM. As suggested by the authors, we kept a mean representation of 1280 dimensions for each mutated sequence. We then reduced the dimensions to 60 using Principal Component Analysis, which was used for downstream supervised learning. The authors’ grid search identified Support Vector Regressor as the best model for variant prediction, which we also used. For leave-oneout cross-validation, we used the Training set described in Methods and Materials. The same workflow was followed for the Prot-T5, using 1024-D embeddings as proposed by the authors.

### Calculation of confidence score

We calculated the average GMM log-probabilities of the amino acids present in each position of the MSA of each protein sequence. We performed a Grid search and selected the confidence formula that optimized the weighted performances across all test sets. This procedure resulted in equation (1) for pathogenic mutations and equation (2) for benign mutations.

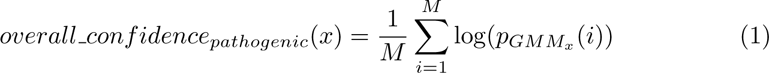

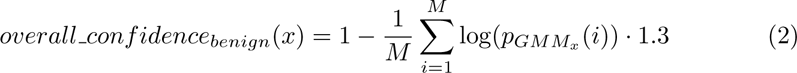

where M the number of samples of fitted GMM (equal to number of sequences in the MSA), log(p*_GMMx_*) the log-probability of each sample under the current model and x the position of the mutation. The inversion of overall confidence for the benign mutations can be interpreted by Supplementary Fig.16 where it is shown that the benign mutations have more confident predictions when the average log-probability of the samples on the position is smaller.

The calculation of the weighted performance was done by multiplying each performance metric by its corresponding weight (i.e overall confidence). This ensures that the contribution of each prediction to the overall performance is proportional to its confidence level. We then summed up the weighted performance metrics for all predictions and divided the sum of the weighted performance metrics by the sum of the weights as shown in equation (3):

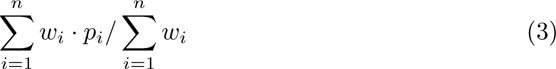

where: *w*_1_*, w*_2_*, . . ., w_n_* are the weights assigned to each prediction and *p*_1_*, p*_2_*, . . ., p_n_* are the performance metrics for each prediction. We proceeded by min-max normalizing the confidence of each prediction to obtain a range of 0-100% for all genes.

##### Key points

- The detection of cancer-driving mutations from genomics data is increasingly feasible, but many challenges remain, particularly detecting such mutations beyond the well-known hotspot regions of cancer-related genes.
- We present the first large-scale benchmark of protein Language Models for predicting cancer-driving mutations.
- Our findings highlight the necessity of including Evolutionary Information in the features captured by protein Language Models, which we achieve with the D2Deep method.
- The D2Deep features capture the epistatic effect of a mutation, which correlates with known mutations in the clinical setting and long-range allosteric effect from other tools
- We introduced a statistical approach to generate confidence scores, crucial for clinical interpretation and mutation prioritization.

## Data availability

All data used for training and testing the model are available in the public Zenodo repository:https://zenodo.org/doi/10.5281/zenodo.8200795 (DOI 10.5281/zenodo.8200795).

## Code availability

All code required to run D2Deep is available via our public github repository: https://github.com/KonstantinaT/D2Deep/.

## Supplementary Figures

**Fig. 7.**
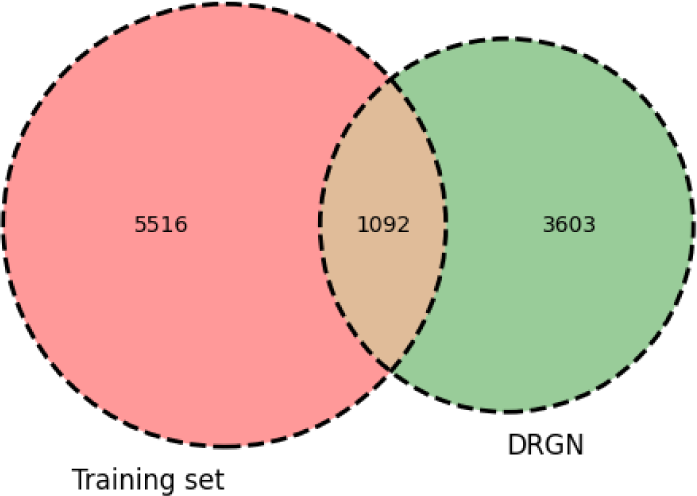
Number of DRGN variants present on Training set.

**Fig. 8.**
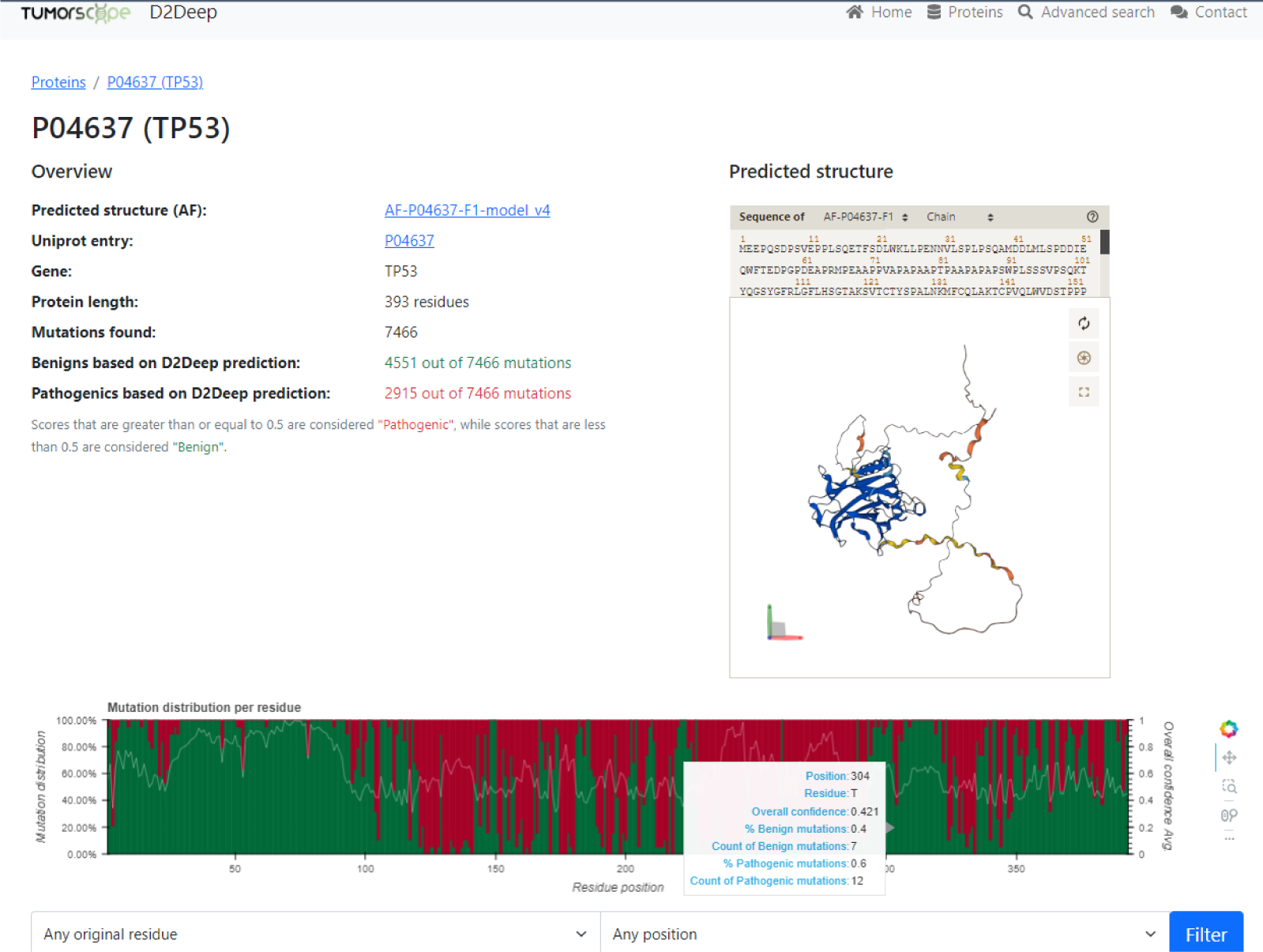
Screenshot of D2Deep’s webserver showing gene view for TP53. An overview of shows the effect of all possible mutations at each position of the gene with the mean confidence score of the predictions. There is also the option of search for specific positions from the option “Advanced search”

**Fig. 9.**
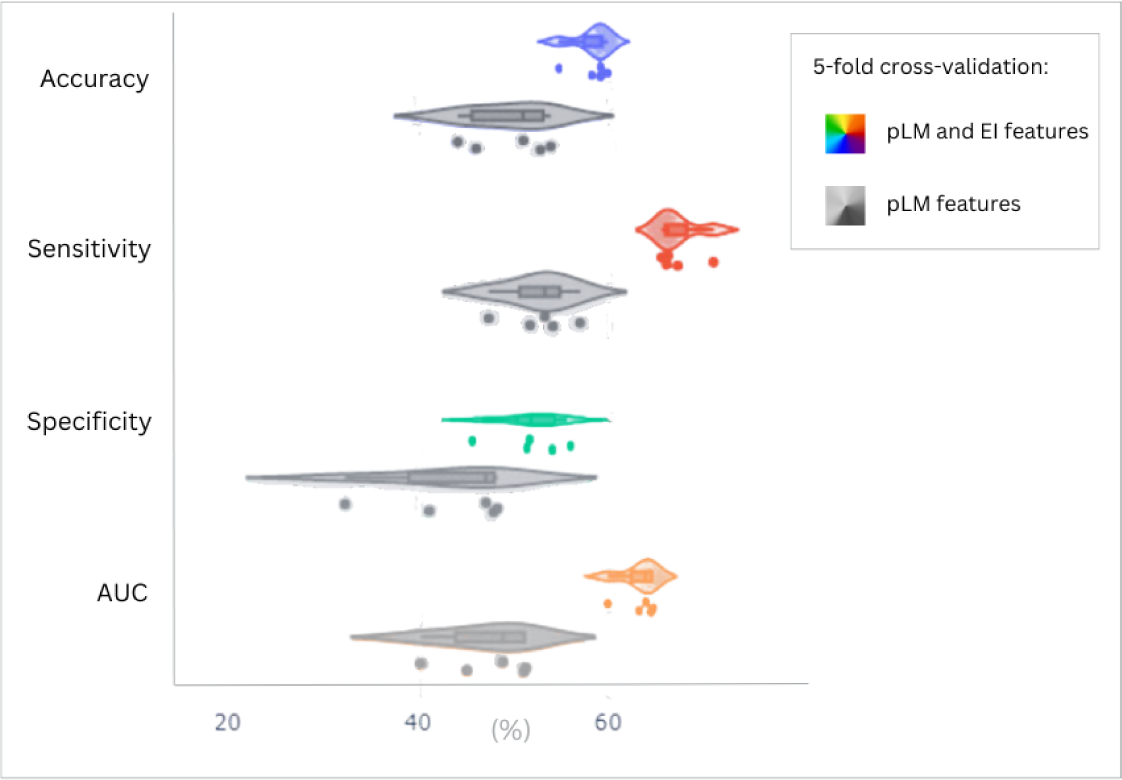
Performance scores running D2Deep in 5-fold cross-validation on DRGN test set when both pLM Information and Evolutionary Information are used (different colors) compared to only pLM Information (grayscale)

**Fig. 10.**
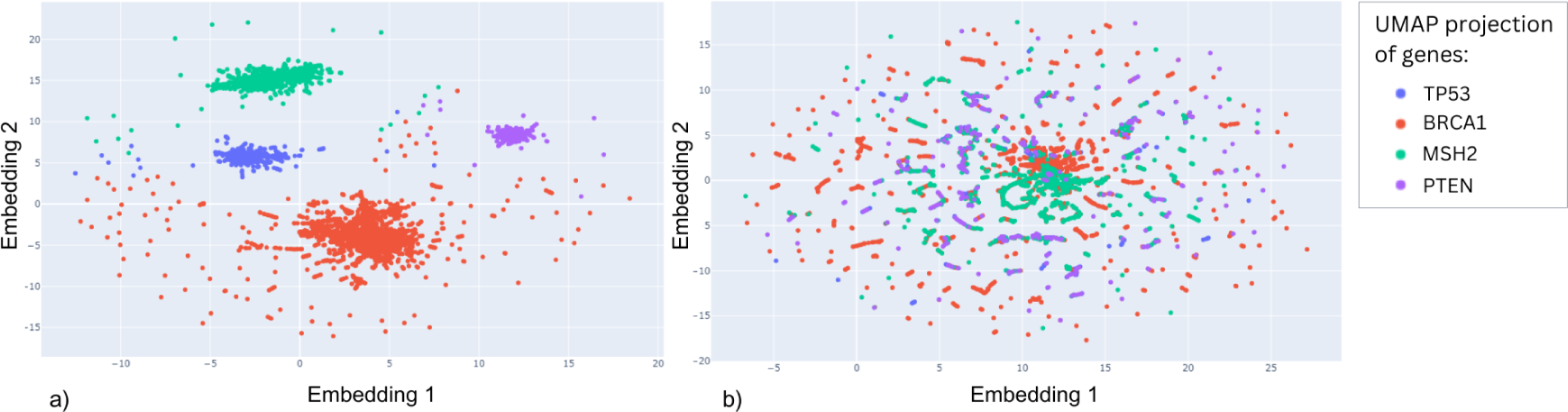
a) We projected in 2D the pLM-based features for the mutated sequences for 4 genes using UMAP dimensionality reduction algorithm. Overall, the generic grammar rules learnt by the model cluster by gene and prevail over the changed induced by the mutation, even though there are regions where amino acid changes in different genes are represented by very similar vectors . b) The UMAP projection of mutated sequences using both Evolutionary Information and pLM features. The mutated sequences are distributed throughout the 2D embedding space

**Fig. 11.**
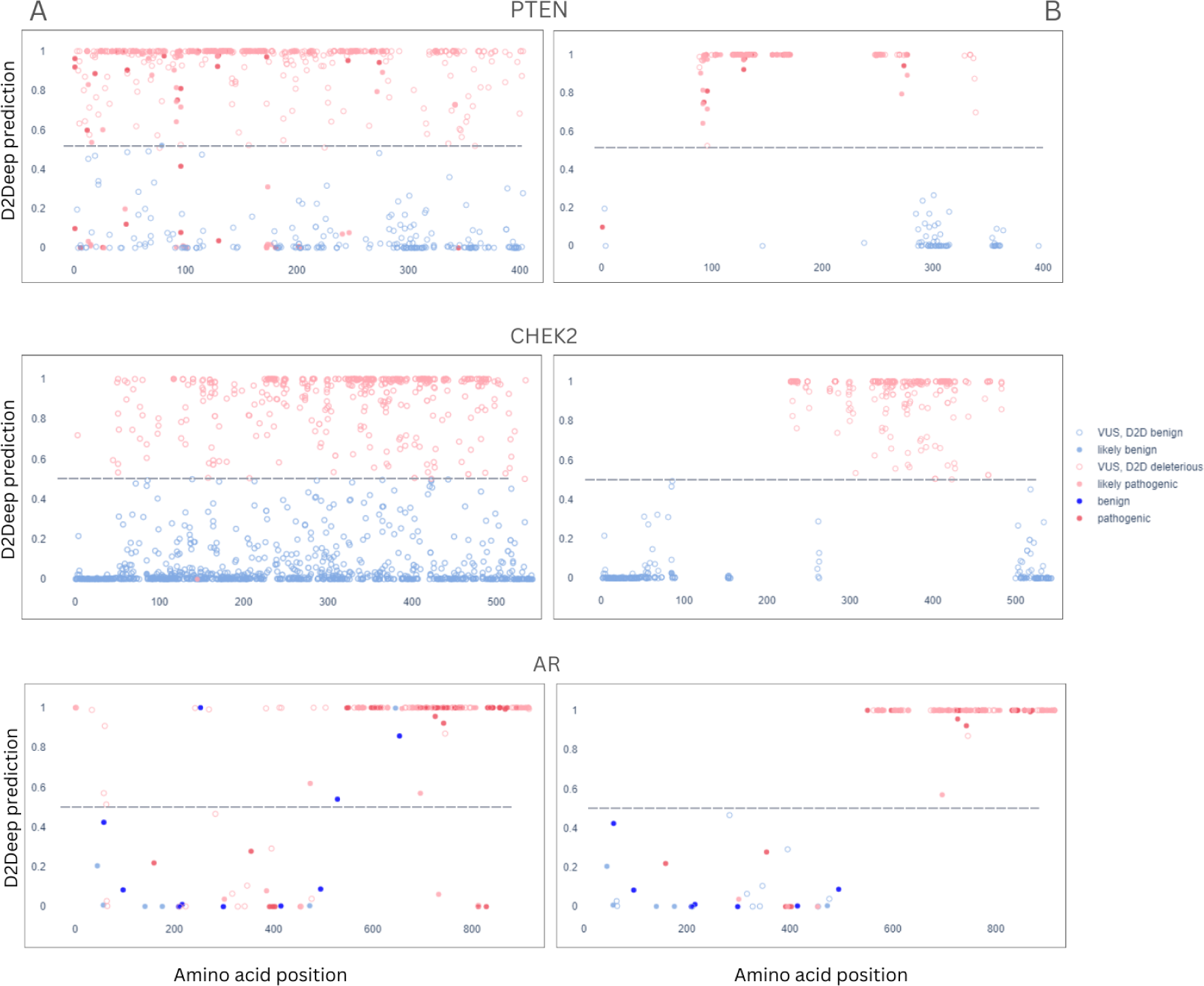
D2Deep results for Clinvar annotations on 3 cancer driver genes (PTEN, CHEK2 and AR) when A: all D2Deeps’ predictions are shown - B: high confidence predictions for Clinvar annotations are shown

**Fig. 12.**
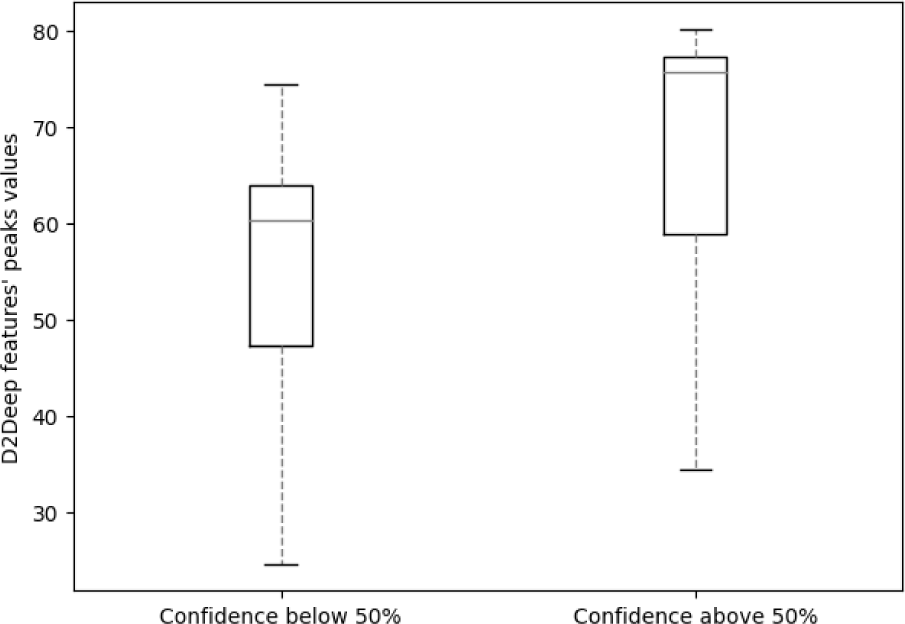
TP53 VUS mutations with confidence score above 0.5 (50%) have bigger peaks than those below 0.5, suggesting that mutated sequences with stronger feature signal have higher confidence of being correctly predicted. As peaks were considered the 10 maximum values of the feature signal used by the predictor.)

**Fig. 13.**
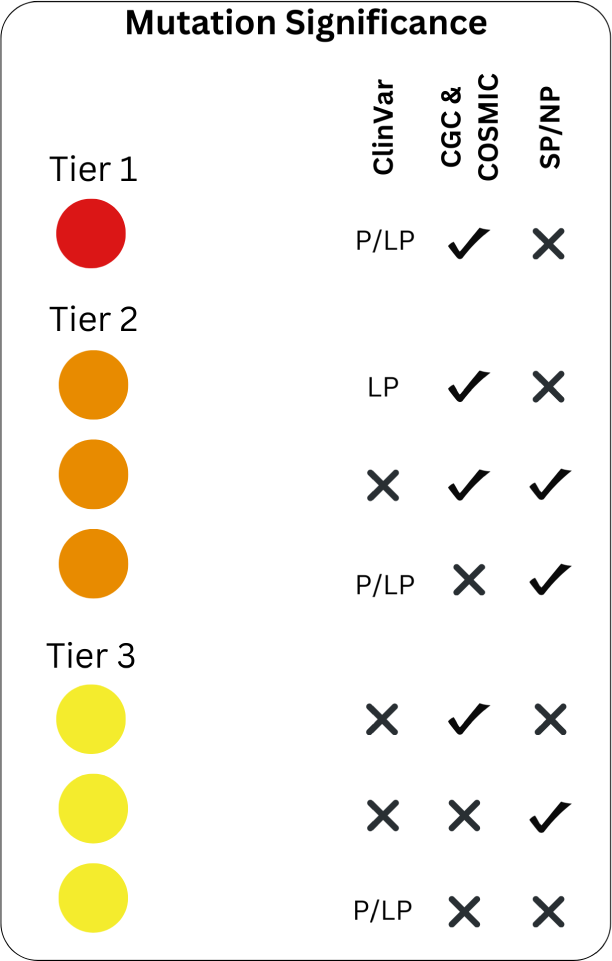
Evidence criteria: 1) Clinvar: classified as pathogenic in Clinvar cancer-related diseases, 2) CGC & COSMIC recurrent missense or small inframe indel in a Cancer Gene Census (CGC) oncogene or Loss-of-function (LOF) frameshift, nonsense, large indel) in a CGC Tumour Suppressor Gene (TSG), 3) SP/NP: Evidence of positive selection as determined by dN/dS algorithm. Reference: https://cancer.sanger.ac.uk/cmc/help.

**Fig. 14.**
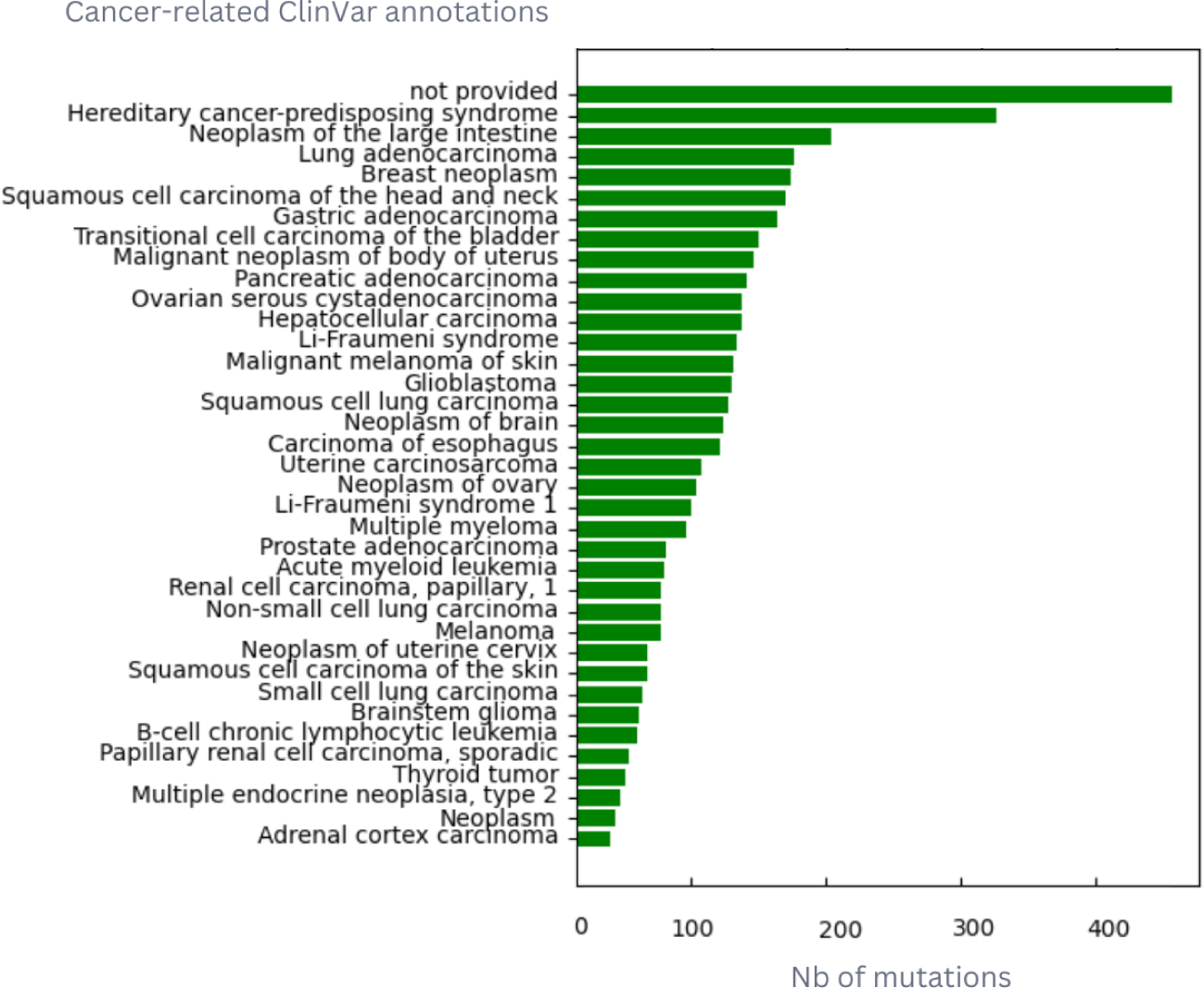
Distribution of Pathogenic or Likely Pathogenic COSMIC mutations in cancer-related diseases in ClinVar (date of download: June 2023).

**Fig. 15.**
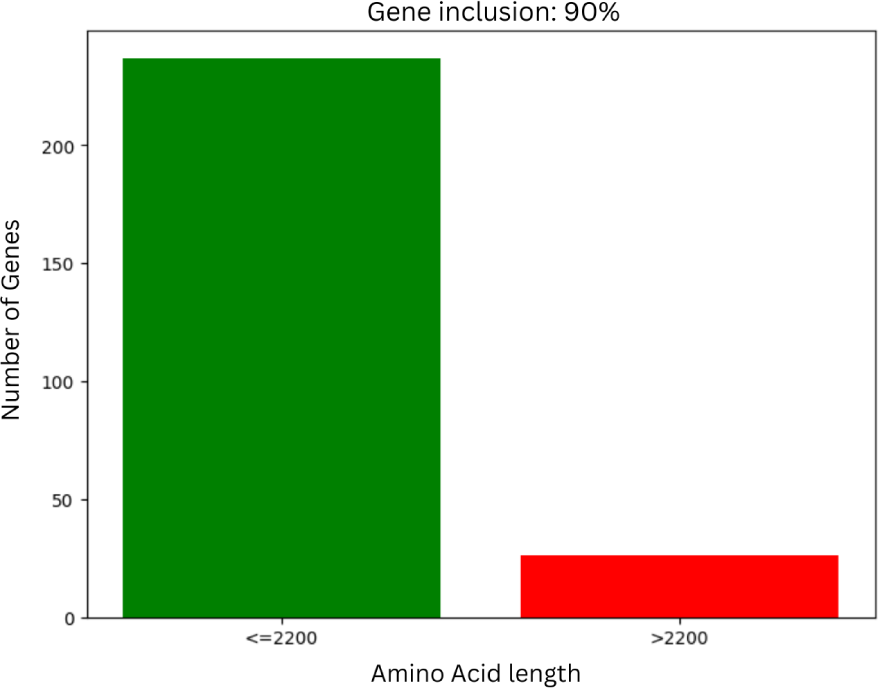
Impact of 2200 Amino Acid length limitation on the number of Compermed genes predicted by D2Deep

**Fig. 16.**
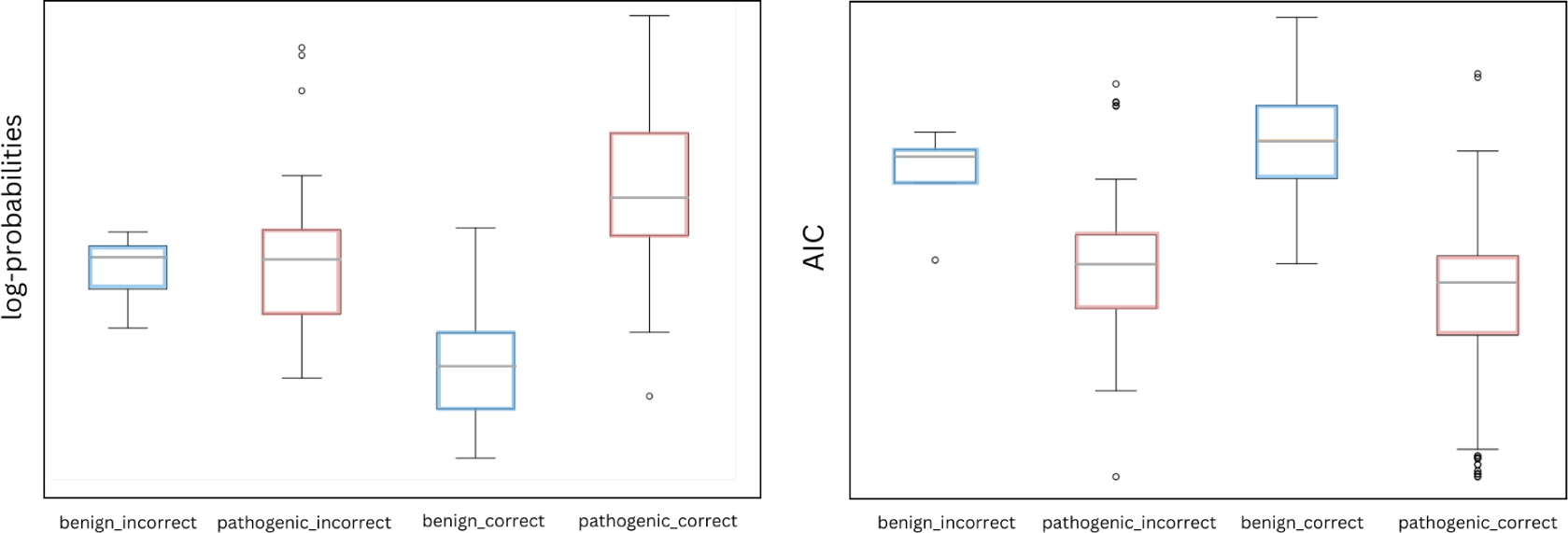
Samples at MSA positions with benign mutations predicted correctly, have smaller GMM log-probability average, while for pathogenic mutations the opposite is observed. Similar conclusions are extracted by the Akaike information criterion (AIC) plot estimating the prediction error - lower values indicate more secure predictions.

**Fig. 17.**
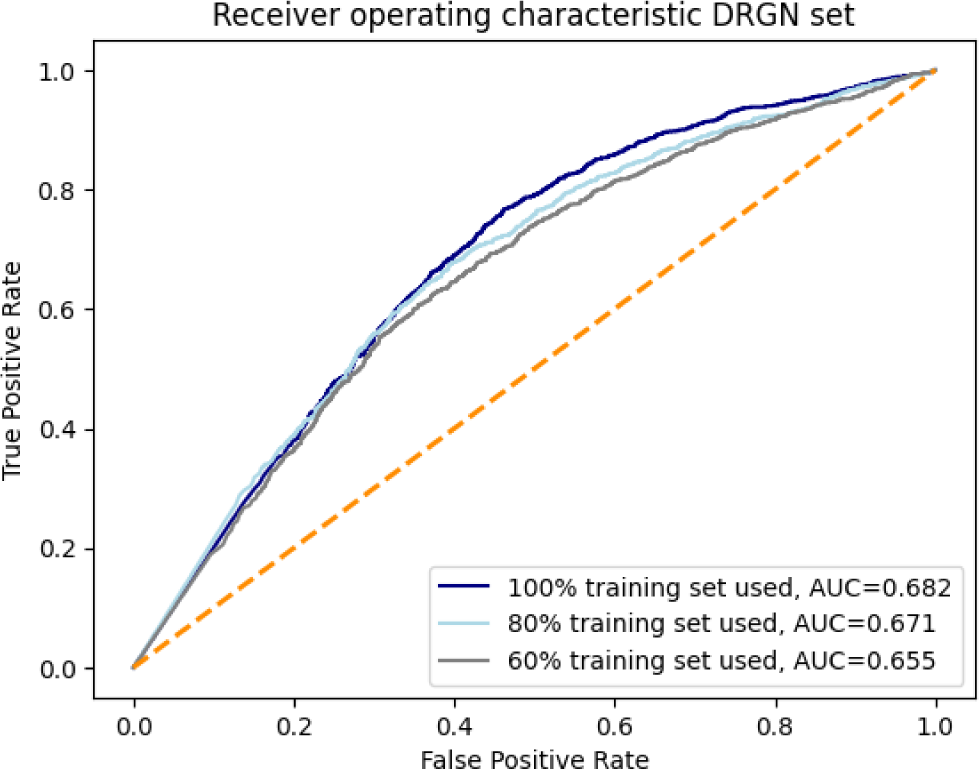
Impact of the size of Training Dataset on the DRGN test set, using 100%, 80% and 60% of the data (Matthews correlation coefficient: 0.31, 0.27 and 0.26 respectively).

**Table 2.** Hyperparameters analysis for DRGN 5-fold cross-validation.

| Predictor's Architecture | Average Sensitivity (%) | Average Specificity (%) | Average AUC(%) |
| --- | --- | --- | --- |
| Presented method (D2D) | **69** | 47.5 | **59.7** |
| Single layer, learning rate 0.0003 | 54.3 | 47.5 | 51.4 |
| Presented method with Dropout 0.1 | 48.1 | 54.1 | 50.3 |
| Single layer, normal initialisation | 61.8 | **75** | 51 |

Examples of hyperparameters tested during Grid Search for the selection of optimal predictive model and their impact on performance on the independent DRGN test set.

## Acknowledgments.

Not applicable

## Declarations

- Funding: Vrije Universiteit Brussel Research Council under the Interdisciplinary Research Program TumorScope [IRP20 to K.T.]; European Union’s Horizon 2020 research and innovation programme under grant agreement No 101016834 (HosmartAI) to K.T., W.V..; Research Foundation Flanders (FWO) International Research Infrastructure [I000323N to W.V.].
- Conflict of interest/Competing interests: Not applicable
- Ethics approval Not applicable
- Consent to participate Not applicable
- Consent for publication Not applicable
- Availability of data and materials: All data used for training and testing the model are available in the public Zenodo repository: https://zenodo.org/records/8200795 (DOI 10.5281/zenodo.8200795).
- Code availability : All code required to run D2Deep is available via our public github repository: https://github.com/KonstantinaT/D2Deep/
- Authors’ contributions K.T developed the computational method and performed data curation and validations. K.T and W.V wrote the manuscript. A.D implemented the webserver. C.O reviewed and edited the manuscript and supervised the research project along with W.V.

## Notes

https://zenodo.org/records/8200795

https://tumorscope.be/d2deep/

https://github.com/KonstantinaT/D2Deep

### Competing Interest Statement

The authors have declared no competing interest.

### Summary of Updates

Methods sections 'Transfer learning with Pre-Trained model ProtT5-XL' and 'Gaussian Mixture Model' updated

## References

[1] Won DG, Kim DW, Woo J, Lee K. 3Cnet: pathogenicity prediction of human variants using multitask learning with evolutionary constraints;37(24):4626–4634. 10.1093/bioinformatics/btab529.

[2] Raimondi D, Passemiers A, Fariselli P, Moreau Y. Current cancer driver variant predictors learn to recognize driver genes instead of functional variants;19(1):3. 10.1186/s12915-020-00930-0.

[3] Jin SC, Homsy J, Zaidi S, Lu Q, Morton S, DePalma SR, et al. Contribution of rare inherited and de novo variants in 2,871 congenital heart disease probands;49(11):1593–1601. Number: 11 Publisher: Nature Publishing Group. 10.1038/ng.3970.

[4] Agarwal SK, Beth Kester M, Debelenko LV, Heppner C, Emmert-Buck MR, Skarulis MC, et al. Germline Mutations of the MEN1 Gene in Familial Multiple Endocrine Neoplasia Type 1 and Related States;6(7):1169–1175. 10.1093/hmg/6.7.1169.

[5] Śevenet N, Sheridan E, Amram D, Schneider P, Handgretinger R, Delattre O. Constitutional mutations of the hSNF5/INI1 gene predispose to a variety of cancers;65(5):1342–1348. 10.1086/302639.

[6] Lamlum H, Al Tassan N, Jaeger E, Frayling I, Sieber O, Reza FB, et al. Germline APC variants in patients with multiple colorectal adenomas, with evidence for the particular importance of E1317Q;9(15):2215–2221. 10.1093/oxfordjournals.hmg.a018912.

[7] Venkitaraman AR. Cancer Susceptibility and the Functions of BRCA1 and BRCA2;108(2):171–182. 10.1016/S0092-8674(02)00615-3.

[8] Cheng J, Novati G, Pan J, Bycroft C, Žemgulytė A, Applebaum T, et al. Accurate proteome-wide missense variant effect prediction with AlphaMissense;381(6664):eadg7492. 10.1126/science.adg7492.

[9] Roy DM, Walsh LA, Chan TA. Driver mutations of cancer epigenomes;5(4):265–296. 10.1007/s13238-014-0031-6.

[10] Greaves M, Maley CC. CLONAL EVOLUTION IN CANCER;481(7381):306–313. 10.1038/nature10762.

[11] Aaltonen LA, Abascal F, Abeshouse A, Aburatani H, Adams DJ, Agrawal N, et al. Pan-cancer analysis of whole genomes;578(7793):82–93. Number: 7793 Publisher: Nature Publishing Group. 10.1038/s41586-020-1969-6.

[12] Ng PC, Henikoff S. SIFT: Predicting amino acid changes that affect protein function;31(13):3812–3814. 10.1093/nar/gkg509.

[13] Adzhubei IA, Schmidt S, Peshkin L, Ramensky VE, Gerasimova A, Bork P, et al. A method and server for predicting damaging missense mutations;7(4):248–249. 10.1038/nmeth0410-248.

[14] Rentzsch P, Witten D, Cooper GM, Shendure J, Kircher M. CADD: predicting the deleteriousness of variants throughout the human genome;47:D886–D894. 10.1093/nar/gky1016.

[15] Shihab HA, Gough J, Cooper DN, Stenson PD, Barker GLA, Edwards KJ, et al. Predicting the Functional, Molecular, and Phenotypic Consequences of Amino Acid Substitutions using Hidden Markov Models;34(1):57–65. 10.1002/humu.22225.

[16] Hopf TA, Ingraham JB, Poelwijk FJ, Schärfe CPI, Springer M, Sander C, et al. Mutation effects predicted from sequence co-variation;35(2):128–135. Number: 2 Publisher: Nature Publishing Group. 10.1038/nbt.3769.

[17] Breen MS, Kemena C, Vlasov PK, Notredame C, Kondrashov FA. Epistasis as the primary factor in molecular evolution;490(7421):535–538. Number: 7421 Publisher: Nature Publishing Group. 10.1038/nature11510.

[18] Figliuzzi M, Jacquier H, Schug A, Tenaillon O, Weigt M. Coevolutionary Landscape Inference and the Context-Dependence of Mutations in Beta-Lactamase TEM-1;33(1):268–280. 10.1093/molbev/msv211.

[19] Pejaver V, Urresti J, Lugo-Martinez J, Pagel KA, Lin GN, Nam HJ, et al. Inferring the molecular and phenotypic impact of amino acid variants with MutPred2;11(1):5918. Number: 1 Publisher: Nature Publishing Group. 10.1038/s41467-020-19669-x.

[20] Weinreich DM, Lan Y, Wylie CS, Heckendorn RB. Should evolutionary geneticists worry about higher-order epistasis?;23(6):700–707. 10.1016/j.gde.2013.10.007.

[21] Domingo J, Diss G, Lehner B. Pairwise and higher order genetic interactions during the evolution of a tRNA;558(7708):117–121. 10.1038/s41586-018-0170-7.

[22] Echave J, Wilke CO. Biophysical models of protein evolution: Understanding the patterns of evolutionary sequence divergence;46:85. Publisher: NIH Public Access. 10.1146/annurev-biophys-070816-033819.

[23] Frazer J, Notin P, Dias M, Gomez A, Min JK, Brock K, et al. Disease variant prediction with deep generative models of evolutionary data;599(7883):91–95. Number: 7883 Publisher: Nature Publishing Group. 10.1038/s41586-021-04043-8.

[24] Brandes N, Goldman G, Wang CH, Ye CJ, Ntranos V. Genome-wide prediction of disease variant effects with a deep protein language model;55(9):1512–1522. Number: 9 Publisher: Nature Publishing Group. 10.1038/s41588-023-01465-0.

[25] Wittmann BJ, Yue Y, Arnold FH. Informed training set design enables efficient machine learning-assisted directed protein evolution;12(11):1026–1045.e7. 10.1016/j.cels.2021.07.008.

[26] Hsu C, Nisonoff H, Fannjiang C, Listgarten J. Learning protein fitness models from evolutionary and assay-labeled data;40(7):1114–1122. 10.1038/s41587-021-01146-5.

[27] Quaio CRD Ceroni JRM, Cervato MC, Thurow HS, Moreira CM, Trindade ACG, et al. Parental segregation study reveals rare benign and likely benign variants in a Brazilian cohort of rare diseases;12:7764. 10.1038/s41598-022-11932-z.

[28] Rives A, Meier J, Sercu T, Goyal S, Lin Z, Liu J, et al. Biological structure and function emerge from scaling unsupervised learning to 250 million protein sequences;p. 46.

[29] Alley EC, Khimulya G, Biswas S, AlQuraishi M, Church GM. Unified rational protein engineering with sequence-based deep representation learning;16(12):1315–1322. Number: 12 Publisher: Nature Publishing Group. 10.1038/s41592-019-0598-1.

[30] Rao R, Meier J, Sercu T, Ovchinnikov S, Rives A.: Transformer protein language models are unsupervised structure learners. bioRxiv. Pages: 2020.12.15.422761 Section: New Results. Available from: https://www.biorxiv.org/content/10.1101/2020.12.15.422761v1.

[31] Elnaggar A, Heinzinger M, Dallago C, Rehawi G, Wang Y, Jones L, et al. Prot- Trans: Toward Understanding the Language of Life Through Self-Supervised Learning;44(10):7112–7127. 10.1109/TPAMI.2021.3095381.

[32] Heinzinger M, Elnaggar A, Wang Y, Dallago C, Nechaev D, Matthes F, et al. Modeling aspects of the language of life through transfer-learning protein sequences;20(1):723. 10.1186/s12859-019-3220-8.

[33] Raimondi D, Tanyalcin I, Ferté J, Gazzo A, Orlando G, Lenaerts T, et al. DEOGEN2: prediction and interactive visualization of single amino acid variant deleteriousness in human proteins;45:W201–W206. 10.1093/nar/gkx390.

[34] Adzhubei I, Jordan DM, Sunyaev SR. Predicting functional effect of human missense mutations using PolyPhen-2;Chapter 7:Unit7.20. 10.1002/0471142905.hg0720s76.

[35] The UniProt Consortium. UniProt: the universal protein knowledgebase;45:D158–D169. 10.1093/nar/gkw1099.

[36] Sonnhammer EL, Eddy SR, Durbin R. Pfam: a comprehensive database of protein domain families based on seed alignments;28(3):405–420. 10.1002/(sici)1097-0134(199707)28:3⟨405::aid-prot10⟩3.0.co;2-l.

[37] Meier J, Rao R, Verkuil R, Liu J, Sercu T, Rives A.: Language models enable zero-shot prediction of the effects of mutations on protein function. bioRxiv. Pages: 2021.07.09.450648 Section: New Results. Available from: https://www.biorxiv.org/content/10.1101/2021.07.09.450648v2.

[38] Dunham AS, Beltrao P. Exploring amino acid functions in a deep mutational landscape;17(7):e10305. 10.15252/msb.202110305.

[39] Arnaudi M, Beltrame L, Degn K, Utichi M, Scrima S, Besora PSI, et al.: MAVISp: A Modular Structure-Based Framework for Genomic Variant Interpretation. bioRxiv. Pages: 2022.10.22.513328 Section: New Results. Available from: https://www.biorxiv.org/content/10.1101/2022.10.22.513328v4.

[40] Garziera M, Cecchin E, Canzonieri V, Sorio R, Giorda G, Scalone S, et al. Identification of Novel Somatic TP53 Mutations in Patients with High-Grade Serous Ovarian Cancer (HGSOC) Using Next-Generation Sequencing (NGS);19(5):1510. 10.3390/ijms19051510.

[41] Saha G, Singh R, Mandal A, Das S, Chattopadhyay E, Panja P, et al. A novel hotspot and rare somatic mutation p.A138V, at TP53 is associated with poor survival of pancreatic ductal and periampullary adenocarcinoma patients;26(1):59. 10.1186/s10020-020-00183-1.

[42] Wang Y, Goh KY, Chen Z, Lee WX, Choy SM, Fong JX, et al. A Novel TP53 Gene Mutation Sustains Non-Small Cell Lung Cancer through Mitophagy;11(22):3587. 10.3390/cells11223587.

[43] Forbes SA, Bhamra G, Bamford S, Dawson E, Kok C, Clements J, et al. The Catalogue of Somatic Mutations in Cancer (COSMIC);Chapter 10:Unit 10.11. 10.1002/0471142905.hg1011s57.

[44] Zhou W, Chen T, Chong Z, Rohrdanz MA, Melott JM, Wakefield C, et al. TransVar: a multilevel variant annotator for precision genomics;12(11):1002–1003. Number: 11 Publisher: Nature Publishing Group. 10.1038/nmeth.3622.

[45] Tamborero D, Rubio-Perez C, Deu-Pons J, Schroeder MP, Vivancos A, Rovira A, et al. Cancer Genome Interpreter annotates the biological and clinical relevance of tumor alterations;10(1):25. 10.1186/s13073-018-0531-8.

[46] Jumper J, Evans R, Pritzel A, Green T, Figurnov M, Ronneberger O, et al. Highly accurate protein structure prediction with AlphaFold;596(7873):583–589. Number: 7873 Publisher: Nature Publishing Group. 10.1038/s41586-021-03819-2.

[47] Suzek BE, Wang Y, Huang H, McGarvey PB, Wu CH, UniProt Consortium. UniRef clusters: a comprehensive and scalable alternative for improving sequence similarity searches;31(6):926–932. 10.1093/bioinformatics/btu739.

